# High-Throughput Screening and CRISPR-Cas9 Modeling of Causal Lipid-Associated Expression Quantitative Trait Locus Variants

**DOI:** 10.1101/056820

**Authors:** Avanthi Raghavan, Xiao Wang, Peter Rogov, Li Wang, Xiaolan Zhang, Tarjei S. Mikkelsen, Kiran Musunuru

## Abstract

Genome-wide association studies have identified a number of novel genetic loci linked to serum cholesterol and triglyceride levels. The causal DNA variants at these loci and the mechanisms by which they influence phenotype and disease risk remain largely unexplored. Expression quantitative trait locus analyses of patient liver and fat biopsies indicate that many lipid-associated variants influence gene expression in a cis-regulatory manner. However, linkage disequilibrium among neighboring SNPs at a genome-wide association study-implicated locus makes it challenging to pinpoint the actual variant underlying an association signal. We used a methodological framework for causal variant discovery that involves high-throughput identification of putative disease-causal loci through a functional reporter-based screen, the massively parallel reporter assay, followed by validation of prioritized variants in genome-edited human pluripotent stem cell models generated with CRISPR-Cas9. We complemented the stem cell models with CRISPR interference experiments *in vitro* and in knock-in mice *in vivo*. We provide validation for two high-priority SNPs, rs2277862 and rs10889356, being causal for lipid-associated expression quantitative trait loci. We also highlight the challenges inherent in modeling common genetic variation with these experimental approaches.

**Author Summary:** Genome-wide association studies have identified numerous loci linked to a variety of clinical phenotypes. It remains a challenge to identify and validate the causal DNA variants in these loci. We describe the use of a high-throughput technique called the massively parallel reporter assay to analyze thousands of candidate causal DNA variants for their potential effects on gene expression. We use a combination of genome editing in human pluripotent stem cells, “CRISPR interference” experiments in other cultured human cell lines, and genetically modified mice to analyze the two highest-priority candidate DNA variants to emerge from the massively parallel reporter assay, and we confirm the relevance of the variants to nearby gene expression. These findings highlight a methodological framework with which to identify and functionally validate causal DNA variants.

## Introduction

Genome-wide association studies (GWASs) have emerged as a powerful unbiased tool to identify single nucleotide polymorphisms (SNPs) associated with incidence of a particular phenotype or disease [1]. Interestingly, only a small fraction of GWAS lead variants lie within coding sequence and thus directly implicate a causal gene at a locus. The vast majority of implicated SNPs fall in noncoding sequence, including introns and gene deserts, suggesting they may play a regulatory role in gene expression. Moreover, most of these SNPs are not themselves causal but exist in linkage disequilibrium (LD) with the true functional variants. The causal gene driving an association signal is often not immediately apparent, unless the locus harbors a gene with a known connection to the phenotype of interest. Although reports of GWASs typically label each associated SNP with the name of the nearest annotated gene or most plausible biological candidate at that locus, experiments in biological models are often necessary to identify the true causal gene at the locus [2–4].

Reassuringly, GWASs for lipid traits have identified variants in loci harboring genes that have previously been implicated in Mendelian disorders of lipoprotein metabolism (i.e. *LDLR*, *PCSK9, ABCA1*, etc). Moreover, GWASs have uncovered a plethora of loci with no prior connection to lipid metabolism. In an extensive GWAS for blood lipids, the Global Lipids Genetics Consortium conducted a meta-analysis of 46 prior lipid GWASs comprising >100,000 individuals of European descent, and identified 95 loci associated with total cholesterol (TC), LDL-C, HDL-C and/or TG [2]. Of these loci, 36 had previously been reported by smaller-scale lipid GWASs at genome-wide significance, while the other 59 were previously unpublished. In principle, these novel loci may offer new insights into lipoprotein metabolism and promising targets for therapeutic intervention.

Although such GWASs have identified a host of associated loci, the pace of functional validation has lagged far behind. For the vast majority of GWAS loci, the causal DNA variants and genes remain unexplored, largely due to the difficult and time-intensive nature of functional follow-up. Even after fine mapping within a locus, as a result of LD tens to hundreds of variants can demonstrate indistinguishably strong associations with the phenotype, suggesting that genetic epidemiology alone is an insufficient means of causal variant discovery.

Because many disease-associated variants are believed to modulate gene expression, expression quantitative trait locus (eQTL) studies may illuminate potential downstream targets of the causal variant. An eQTL is a genomic region that influences gene expression either in *cis* or *trans*; however, GWAS-implicated variants are predominantly believed to function in *cis*-acting manner. Importantly, eQTL studies have identified differentially regulated transcripts hundreds of kilobases away from the genotyped variant, implicating long-range chromatin looping interactions in the mechanisms of some eQTLs [5,6]. These differentially regulated genes then become candidates for experimental manipulation (for example, through overexpression and knockout of the orthologous genes in murine models) to ascertain their relevance to the phenotype of interest [2]. However, identifying the causal variants underlying eQTLs through functional studies is more challenging due to the poor conservation of noncoding DNA across species.

Further insight into eQTL mechanisms can be gleaned by integrating risk-associated variants with annotated maps of regulatory elements, such as DNase I hypersensitivity sites, chromatin immunoprecipitation sequencing (ChIP-seq) peaks, and histone modifications. Candidate SNPs that fall in transcriptionally active regions can then be prioritized for functional investigation. Experimental approaches such as reporter assays, electrophoretic mobility shift assays (EMSA), and ChIP can be employed to investigate allele-specific regulatory activity at each variant, as well as determinants of differential transcription factor binding and function [7–9]. However, such efforts have been laborious, requiring a significant amount of dedicated effort to functionally validate each candidate SNP.

Two recently emergent technologies make it feasible to interrogate risk-associated variants in eQTL loci in a much higher throughput fashion. The massively parallel reporter assay (MPRA) allows investigators to generate high-complexity pools of reporter constructs where each regulatory element or variant of interest is linked to a synthetic reporter gene that carries an identifying barcode [10,11]. The reporter construct pools are introduced into relevant populations of cultured cells, and the relative transcriptional activities of the individual elements or variants are measured by sequencing the transcribed reporter mRNAs and counting their specific barcodes. This approach can be used to rapidly profile the regulatory activity of thousands of variants linked to eQTLs for specific phenotypes of interest.

Advances in genome editing technologies—from first-generation zinc finger nucleases (ZFNs) to, more recently, transcription activator-like effector nucleases (TALENs) and clustered regularly interspaced short palindromic repeats (CRISPR)-CRISPR-associated 9 (Cas9) systems—have opened up unprecedented avenues by which to rigorously assess the functional impact of novel genetic variants [12]. All three of these genome-editing tools can be used to introduce targeted alterations into mammalian cells and model organisms. However, CRISPR-Cas9 offers an optimal combination of high targeting efficiency, ease of use, and scalability [13]. The most widely used CRISPR-Cas9 system employs the *Streptococcus pyogenes* Cas9 nuclease, which complexes with a synthetic guide RNA (gRNA) encoding a site-specific 20-nt protospacer sequence that hybridizes an N20NGG target DNA sequence. Once Cas9 induces a double-strand break three nucleotides upstream of the NGG sequence, or protospacer adjacent motif (PAM), the cell employs error-prone non-homologous end joining (NHEJ) to repair the break, often leading to the introduction of an insertion or deletion that may disrupt gene function. If a singlestrand oligonucleotide (ssODN) or double-strand DNA vector is introduced, the cell can utilize it as a donor template for homology-directed repair (HDR), enabling knock-in of specific mutations.

With a goal of better understanding the role of human genetic variation in influencing blood lipid levels, we employed a combination of MPRAs and CRISPR-Cas9 genome editing to high-throughput screen, identify, and validate causal variants at eQTL loci that have been linked to blood lipids in humans.

## Results

### Massively parallel reporter assays define two lipid-associated sites with significant transcriptional activity

The Global Lipids Genetics Consortium interrogated lead SNPs at 95 loci for blood lipids against transcript abundance of local genes in biopsy samples of human liver, subcutaneous fat, and omental fat [2]. This analysis identified 57 total eQTLs, suggesting that many lipid-associated causal SNPs influence gene expression in a cis-regulatory manner. These eQTLs may highlight candidate lipid-modulating genes, possibly located up to hundreds of kilobases away from the eQTL tag SNPs, that underlie the GWAS association signals at these loci. However, LD among neighboring SNPs at any given eQTL locus makes it challenging to pinpoint the causal variant underlying an association signal.

To address this issue, we performed an MPRA experiment to rapidly profile the regulatory activity of 1,837 variants linked to the 16 eQTL lead SNPs identified for subcutaneous fat and/or omental fat (**S1 Table**; see Supplementary Tables 9 and 10 in ref. 2 for more information on eQTLs) and thus prioritize potentially causal candidate SNPs. We used a pool of reporter constructs in which every plausible regulatory variant (i.e., all SNPs with *r*^2^ ≥ 0.5 relative to the 16 eQTL lead SNPs) was embedded within a 144-bp genomic “tile” in six versions—major or minor allele in the center, towards the 5’ end (right-shifted), or towards the 3’ end (left-shifted). Each fragment was coupled to a reporter gene with a unique barcode identifier in the 3’ untranslated region. To minimize barcode-and amplification-associated biases, each fragment was coupled to ~22 distinct barcodes. The pool was transfected into cultured mouse 3T3-L1 adipocytes or pre-adipocytes, and the copy number of each barcode was determined by RNAseq and normalized to the amount of corresponding reporter DNA plasmid that entered the cells [10]. From the MPRA data (**S2 Table**), we prioritized variants that displayed significant allele-specific enhancer activity, as measured by reporter expression, in 3T3-L1 adipocytes. We selected the two variants with the strongest evidence, rs2277862 and rs10889356, for further investigation (**Fig 1A**).

**Fig. 1.**
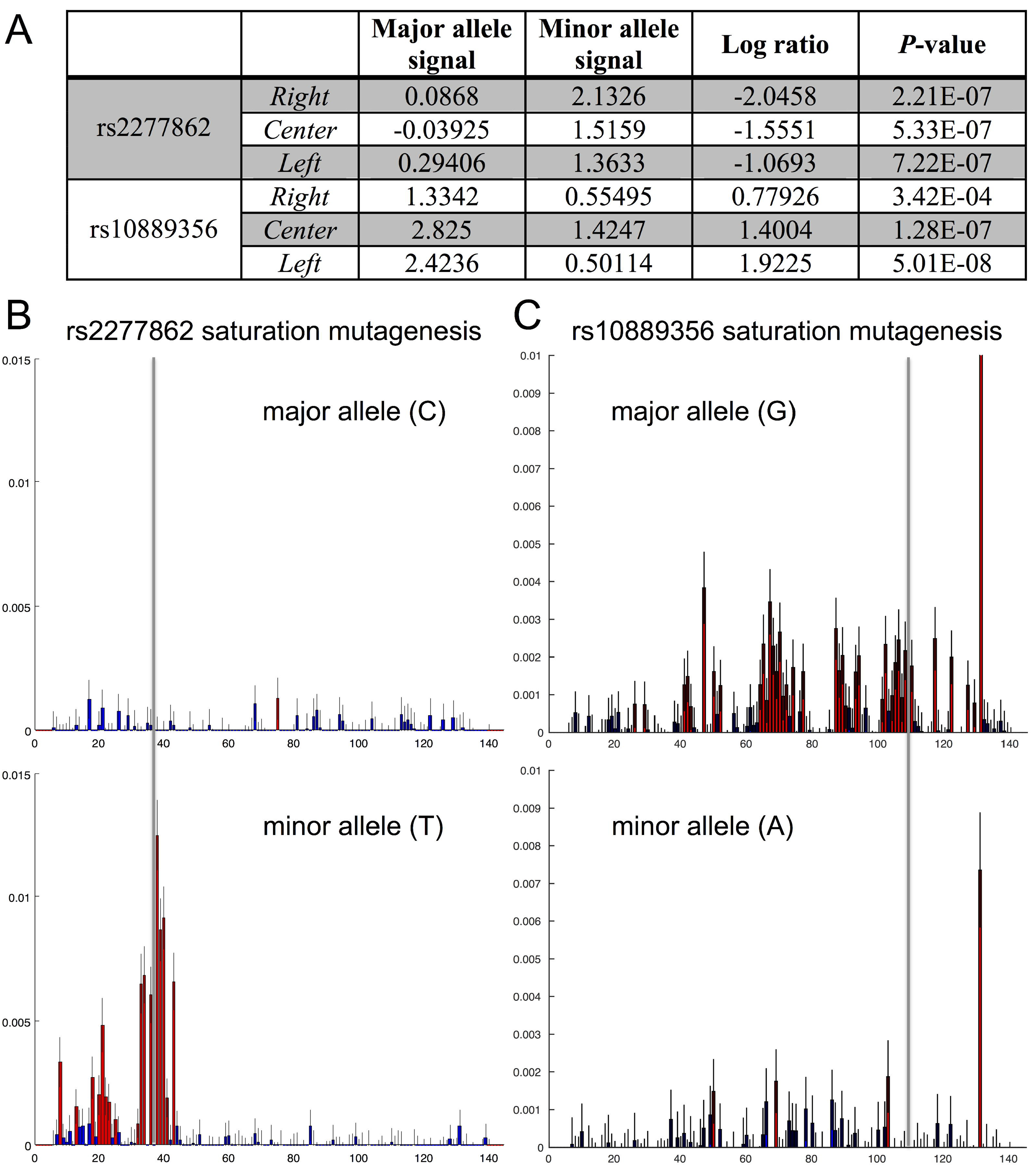
Results of massively parallel reporter assays. (A) MPRA identified rs2277862 and rs10889356 as the SNPs with highest allele-specific regulatory activity in mouse 3T3-L1 adipocytes. For this experiment, each candidate SNP was represented on a 144-bp tile that was either centered, left-shifted, or right-shifted relative to the SNP, in order to increase the probability of capturing the correct regulatory context for that SNP. For each tile, the individual signals for the major and minor alleles are shown for a representative experiment (where signal refers to the log of median barcode counts for the given tile divided by median barcode counts for all tiles). A positive signal implies enhancer activity, while a negative signal implies repressor activity. In the final two columns, a log-ratio for major allele signal over minor allele signal is calculated, along with a-value (by Mann-Whitney *U* test) for the null hypothesis that the major and minor alleles generate equal signals. (B) MPRA-based single-hit saturation mutagenesis experiment with the rs2277862 right-shifted tile, either with the major allele of the SNP (top) or the minor allele (bottom). Red bars indicate a significant change from the original tile’s activity (Mann-Whitney *U* test, 5% FDR); blue bars, not significant. (C) MPRA-based single-hit saturation mutagenesis experiment with the rs10889356 left-shifted tile, either with the major allele of the SNP (top) or the minor allele (bottom).

The Global Lipids Genetics Consortium found rs2277862 to be the lead SNP for total cholesterol (TC) at chromosome 20q11 (*P* = 4 × 10^−10^), with the minor allele associated with a 1.19 mg/dL decrease in TC [2]. This locus harbors several genes with no prior connection to lipid metabolism, including *ERGIC3, CPNE1*, and *CEP250*. rs10889356 is tightly linked to the lead SNP for triglycerides (TG) (*P* = 9 × 10^−43^), low-density-lipoprotein cholesterol (LDL-C), and TC at chromosome 1p31, which harbors *DOCK7*and *ANGPTL3*, the latter a well-established modulator of blood lipids [14]. We hypothesized that rs2277862 and rs10889356 are causal for their respective eQTLs and that each lies within a transcriptional regulatory site that influences local gene expression in an allele-specific manner.

To better define the transcriptional activity at the sites of each of the two SNPs, we performed single-hit saturation mutagenesis experiments with MPRAs in 3T3-L1 adipocytes using pools of reporter constructs harboring 144-bp tiles around each SNP with mutagenesis at each position in the tiles. This enabled us to generate “footprint” representations (**Figs 1B** and **1C**) for each site [10]. For rs2277862, when the minor allele (T) was present, there was a signal indicative of a factor binding directly to the site of the SNP (CAAATA[T]GGCGA) and another signal indicative of a second factor binding upstream of the first factor (ACGAGGTCA), with no signal observed downstream of the site of the SNP (**Fig 1B**). When the major allele (C) was present, all signal disappeared, suggesting that this allele abolishes all of the binding interactions. For rs10889356, when the major allele (G) was present, there was a signal indicative of a factor binding directly to the site of the SNP (AACTTCCT[G]T), as well as several other signals indicative of multiple other factors binding on either side of the first factor (**Fig 1C**). When the minor allele (A) was present, almost all signal disappeared. Of note, there was one downstream nucleotide that showed a very strong signal regardless of allelic variation at rs10889356, suggesting it contributes to transcriptional regulation in a manner independent of the SNP. These data suggest that, for both rs2277862 and rs10889356, complexes of factors that are in part anchored at the site of the SNP are responsible for modulation of local gene expression.

### Genome editing of the rs2277862 site in human pluripotent stem cells

The MPRA data from 3T3-L1 adipocytes indicate that the minor allele (T) of rs2277862 (minor allele frequency in Europeans = 0.15) increases transcriptional activity relative to the major allele (C) (**Fig 1**). This would suggest that the minor allele functions as an enhancer, the major allele functions as a repressor, or both. The rs2277862 locus harbors a number of genes with no prior connection to lipid metabolism, although only three—*CEP250, CPNE1* and *ERGIC3*—show evidence of differential regulation in human liver, subcutaneous fat, and omental fat biopsies. The biology and relevant sites of action of all three of these genes are poorly understood. Interestingly, rs2277862 is located over 50 kb away from two of these eQTL genes, *CEP250* and *CPNE1*, suggesting that it may function as a long-range enhancer or repressor (**Fig 2A**).

**Fig. 2.**
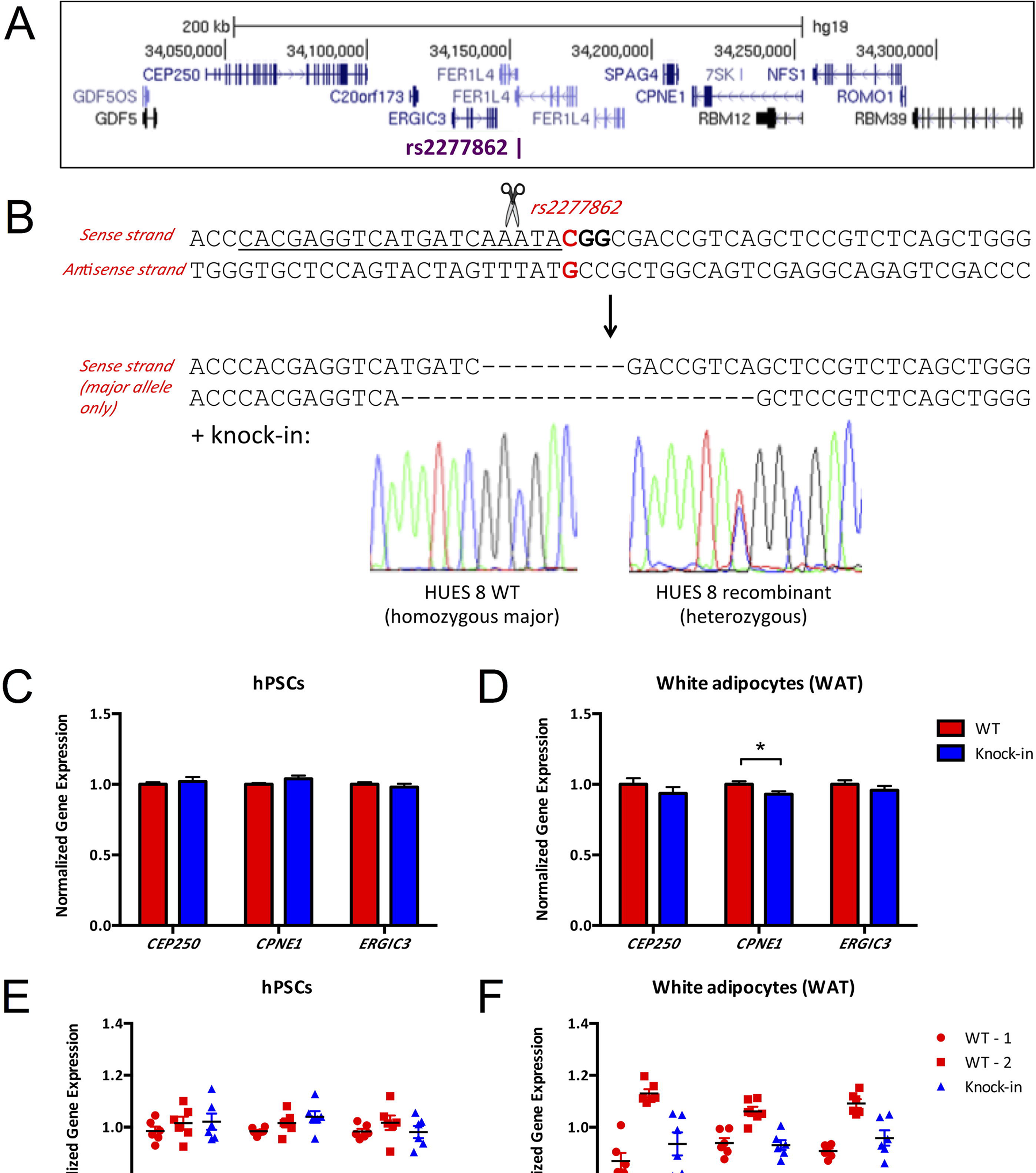
CRISPR-Cas9 genome editing at the rs2277862 locus in hPSCs. (A) Schematic of human chromosome 20q11 locus showing the relative positions of rs2277862, *CEP250, CPNE1*, and *ERG1C3*. (B) Heterozygous rs2277862 minor allele knock-in was generated on the HUES 8 background (homozygous major at rs2277862) at 0.15% frequency using a gRNA that cuts 3 bp upstream from the SNP and an exogenous ssODN. Representative indels are also shown. The guide RNA protospacer is underlined, the PAM is bolded, and the SNP position is indicated in red. (C) Gene expression in undifferentiated HUES 8 cells (*N* = 2 wild-type clones and 1 knock-in clone; 6 wells per clone), normalized to mean expression level in wild-type clones. (D) Gene expression in differentiated HUES 8 adipocytes (*N* = 2 wild-type clones and 1 knock-in clone; 6 wells per clone) (E, F) Individual data points for each clone shown in (C) and (D). Data are displayed as means and s.e.m. (^*^P<0.05, ^**^P<0.01, ^***^P<0.001 relative to control, calculated by two-tailed unpaired Student’s *t*-test).

To define the contribution of each allele to gene expression, we sought to alter the putative regulatory element encompassing rs2277862 in two hPSC lines with different genotypes at this SNP: HUES 8 (C/C, major/major) and H7 (T/T, minor/minor). Although the precise regulatory elements at this locus remain to be determined, the saturation mutagenesis MPRA data for the site suggests that rs2277862 lies directly within such an element. We began by attempting to knock in the minor allele of rs2277862 (T) onto the HUES 8 (C/C) background via HDR. Using CRISPR-Cas9 with a single gRNA with a predicted cleavage site 3 bp upstream from the SNP along with a 67-nt ssODN in HUES 8 cells, we obtained a single recombinant heterozygote at a frequency of 0.15% (1 out of 672 clones screened) (**Fig 2B**).

Because rs2277862 has an eQTL in three developmentally distinct tissues—liver, subcutaneous fat, and omental fat—we reasoned that it may function as a global, non-cell-type-restricted regulator of gene expression and thus modulate transcription in undifferentiated hPSCs as well. We analyzed gene expression and observed no difference in *CEP250, CPNE1*, or *ERGIC3* expression between wild-type clones (C/C) and the recombinant heterozygote clone (C/T) (*n* = 2 wild-type clones and 1 knock-in clone; 6 wells per clone) (**Fig 2C**). We then differentiated the three clones into adipocytes, the cell type in which the MPRA was originally performed. We reasoned that differentiated adipocytes might have greater transcriptional activity at the rs2277862 locus, perhaps by recruitment of additional transcriptional machinery to the site, and thereby exhibit more pronounced gene expression differences. However, hPSC-derived knock-in adipocytes (*n* = 2 wild-type clones and 1 knock-in clone; 6 wells per clone) demonstrated altered expression of only one of the 20q11 genes, *CPNE1* (down 6.9%) (**Fig 2D**).

Analysis of the data revealed that the differentiation protocol had introduced a great deal of stochastic variation in gene expression, not only among genetically identical clones but also among independent wells of the same clone. This point is emphasized in **Figs 2E** and **2F**, where we have plotted the individual data points for the clones analyzed in **Figs 2C** and **2D** (with each point representing a different well of the indicated clone). hPSCs demonstrated minimal intra-clonal well-to-well variability, as well as minimal inter-clonal variability, based on comparison of the two wild-type clones. However, when the same clones were differentiated to adipocytes, the two wild-type clones displayed fairly distinct “setpoints” in gene expression, presumably due to variations in differentiation efficiency. Thus, this variability between genetically equivalent clones within a group could be confounding the ability to discern true differences secondary to a genetic modification.

In light of the inefficiency of the HDR knock-in at the rs2277862 locus, we adopted an alternative approach. Reasoning that small deletions encompassing the site of the SNP should have effects on gene expression, since transcription factor binding sites are typically 8-10 nucleotides long, we used CRISPR-Cas9 to generate numerous deletion mutants. One disadvantage of using NHEJ to introduce indels at a genomic site with a single gRNA is that a wide variety of insertions and deletions are generated, and different indels may have different effects on transcriptional regulatory activity. We therefore utilized a different strategy with two gRNAs with cleavage sites flanking the SNP, 38 bp apart. Through this multiplexing NHEJ approach we efficiently generated small deletions encompassing the candidate SNP at a frequency of 59% (168 out of 285 clones screened) (**Fig 3A**). The use of dual gRNAs facilitated efficient generation of many hPSC clones harboring predictable, and often homozygous, deletions. Due to the efficacy of the multiplexing strategy, we were easily able to generate a large number of homozygous deletion (“knockout”) cell clones in both the HUES 8 (C/C) and H7 (T/T) cell lines.

**Fig. 3.**
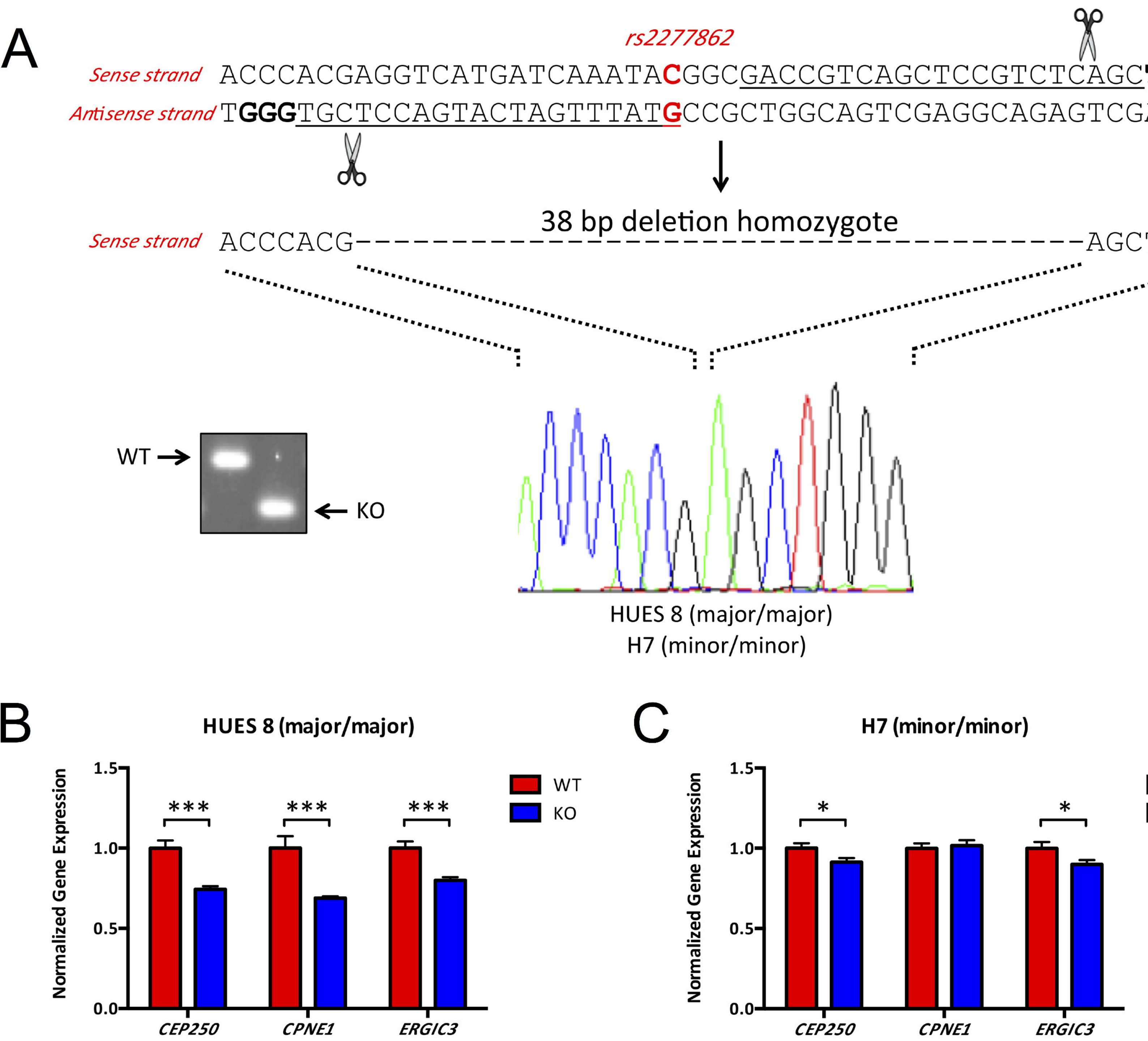
Genetic deletion of rs2277862 site in hPSCs alters gene expression at the 20q11 locus. (A) Homozygous 38-bp deletions (“knockout”) encompassing rs2277862 were generated on the HUES 8 (homozygous major) and H7 (homozygous minor) backgrounds using a dual gRNA approach. A representative agarose gel of PCR amplicons is shown. The guide RNA protospacers are underlined, the PAMs are bolded, and the SNP position is indicated in red. (B) Gene expression in undifferentiated HUES 8 cells (*N* = 10 wild-type and 10 knockout clones, 3 wells per clone), normalized to mean expression level in wild-type clones. (C) Gene expression in undifferentiated H7 cells (*N* = 8 wild-type and 6 knockout clones, 3 wells per clone). Data are displayed as means and s.e.m. (^*^P<0.05, ^**^*P*<0.01, ^***^*P*<0.001 relative to control, calculated by two-tailed unpaired Student’s t-test).

By quantitative reverse transcriptase-polymerase chain reaction (qRT-PCR), we observed statistically significant changes in the expression of 20q11 genes in both hPSC lines. Homozygous disruption of the major allele in HUES 8 cells (*n* = 10 wild-type and 10 knockout clones, 3 wells per clone) substantially decreased expression of *CEP250* (down 25.7%), *CPNE1* (down 31.2%) and *ERGIC3* (down 20.0%) (**Fig 3B**). Homozygous disruption of the minor allele in H7 cells (*n* = 8 wild-type and 6 knockout clones, 3 wells per clone) subtly decreased expression of *CEP250* (down 8.8%) and *ERGIC3* (down 10.2%) (**Fig 3C**).

In combination, the data from genome editing of the rs2277862 site in hPSCs suggest that the major allele of rs2277862 functions as a transcriptional enhancer in hPSCs, and the minor allele may have minimal enhancer activity as well. Intriguingly, these data are at odds with the MPRA data, which suggest that the minor allele increases transcriptional activity relative to the major allele. Nonetheless, the MPRA and genome-editing data agree that rs2277862 is a causal variant with respect to transcriptional regulation.

### Genome editing of the rs10889356 site in human pluripotent stem cells

Rs10889356 is tightly linked to rs2131925 (*r*^2^ = 0.90), the lead SNP for TG, TC and LDL-C at the 1p31 locus, and possession of the minor allele at rs2131925 (MAF = 0.32) is associated with a 4.9 mg/dL decrease in TG [2]. rs10889356 is situated in the promoter of the *DOCK7* gene, which encodes a guanine nucleotide exchange factor that has not previously been implicated in lipid metabolism. *ANGPTL3*, the probable causal gene at this locus, lies within an intron of *DOCK7* and encodes a liver-specific secreted protein that inhibits endothelial lipase and lipoprotein lipase, thereby increasing circulating levels of TG and HDL-C (**Fig 4A**) [14]. Curiously, eQTL data for rs2131925 suggest that expression levels of *DOCK7* and *ANGPTL3* are inversely related in human liver samples, implying that the causal variant variably upregulates or downregulates transcription of different genes at this locus through an unknown mechanism [2].

**Fig. 4.**
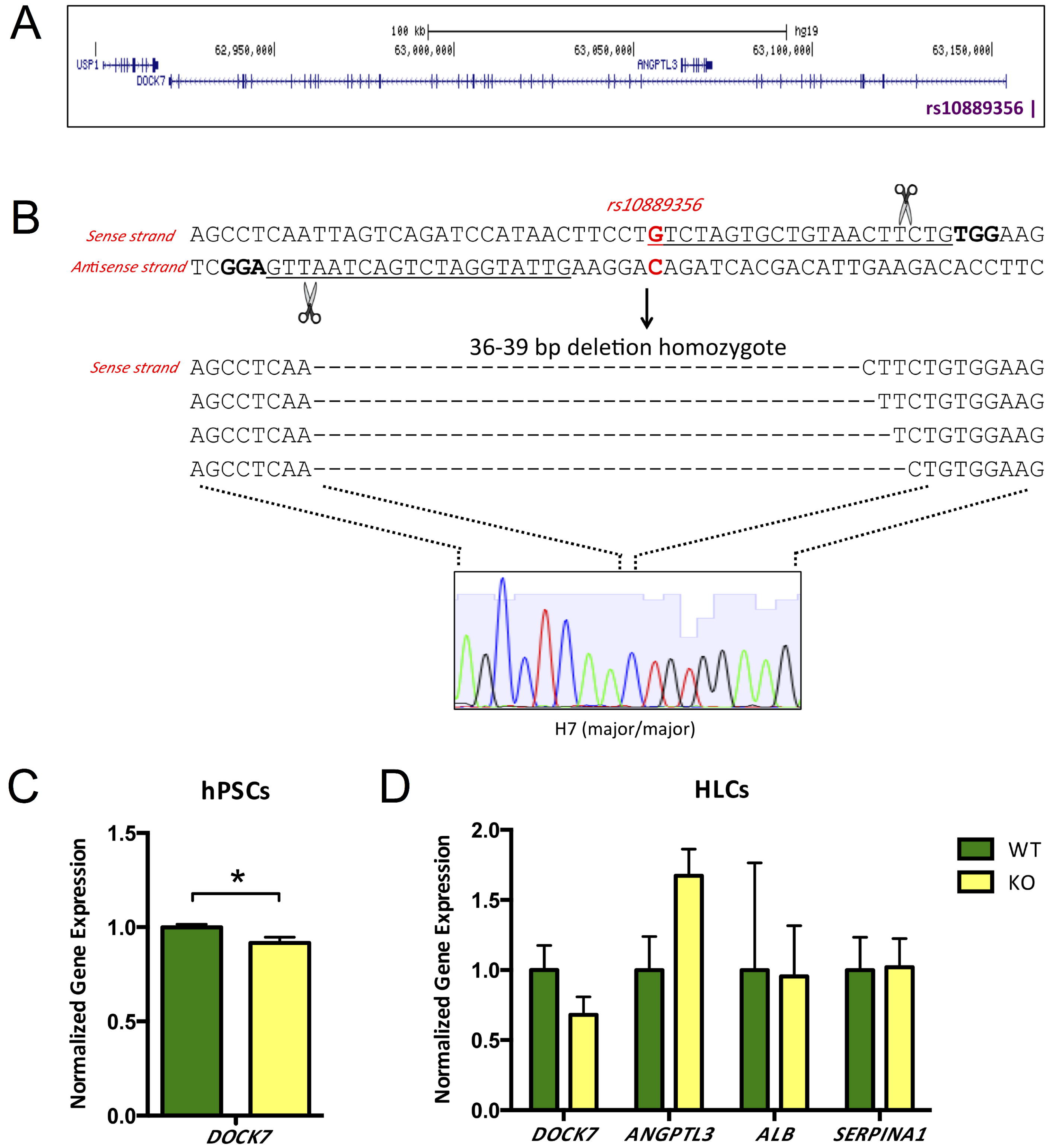
Genetic deletion of rs10889356 site in hPSCs alters gene expression at the 1p31 locus. (A) Schematic of human chromosome 1p31 locus showing the relative positions of rs10889356, *DOCK7*, and *ANGPTL3*. (B) Homozygous 36-to 39-bp deletions (“knockout”) encompassing rs10889356 were generated on the H7 (homozygous major) background using a dual gRNA approach. The guide RNA protospacers are underlined, the PAMs are bolded, and the SNP position is indicated in red. (C) *DOCK7* expression in undifferentiated H7 cells (*N* = 12 wild-type and 8 knockout clones, 3 wells per clone), normalized to mean expression level in wild-type clones. (D) Gene expression in differentiated H7 hepatocyte-like cells (*N* = 4 wild-type and 4 knockout clones, 6 wells per clone). Data are displayed as means and s.e.m. (^*^*P*<0.05, ^**^*P*<0.01, ^***^*P*<0.001 relative to control, calculated by two-tailed unpaired Student’s *t*-test).

The MPRA data indicate that the major allele (G) of rs10889356 has enhancer activity relative to the minor allele (A). Using dual gRNAs, we efficiently generated homozygous deletion mutants for rs10889356 in H7 cells, which are homozygous major (G/G) at this SNP. We observed some heterogeneity in deletion size, presumably because one of the gRNAs did not always induce a DSB exactly at the predicted location 3 bp upstream from the PAM (**Fig 4B**). For gene expression studies, we utilized hPSC clones harboring a range of 36- to 39-bp homozygous deletions as we were not able to obtain a sizeable number of deletion mutants of one particular genotype. In undifferentiated hPSCs (*n* = 12 wild-type and 8 knockout clones, 3 wells per clone), disruption of the major allele diminished expression of *DOCK7* (down 8.3%), suggesting that the major allele indeed confers enhancer activity (**Fig 4C**). *ANGPTL3* expression was too low to be reliably measured in undifferentiated hPSCs, consistent with its being a liver-specific gene.

We then differentiated a subset of clones (*n* = 4 wild-type and 4 knockout clones, 6 wells per clone) to hepatocyte-like cells (HLCs). The HLCs were characterized by expression of the liver-specific markers *ALB* and *SERPINA1*, and the average levels of both these transcripts were roughly equivalent between wild-type and knockout clones. Expression of *DOCK7* was decreased in knockout HLCs, while expression of the liver-specific gene *ANGPTL3* was increased (**Fig 4D**). However, these differences did not achieve statistical significance due to marked variation in differentiation efficiency, not only among clones but also among different wells from the same clone. Indeed, while average *ALB* expression levels were equivalent between groups, the dynamic range of *ALB* expression within each group exceeded an order of magnitude (data not shown), highlighting the tremendous variability of the HLC differentiation protocol, paralleling our findings with the adipocyte differentiation protocol used in studying the rs2277862 heterozygous knock-in clone.

### CRISPR interference at rs2277862 and rs10889356 sites

One possible complementary approach to validating a candidate causal SNP is to harness the sequence-dependent targeting specificity of CRISPR-Cas9 to direct a catalytically dead Cas9 mutant (dCas9) to the SNP site. The rationale is that if a variant is truly causal and lies within a transcriptional regulatory element, then the presence of the bulky dCas9 protein at the site could sterically hinder recruitment of native transcription factors to the regulatory site and thus interfere with transcriptional regulation, i.e., CRISPR interference (CRISPRi) [15].

We generated CRISPRi constructs that co-expressed dCas9 with enhanced green fluorescent protein (EGFP) and expressed each of the three gRNAs shown in **Fig 5A**. We transiently transfected HEK 293T cells, which are homozygous for the major allele (C/C) at rs2277862, with the dCas9 and gRNA constructs, either singly or in combination. With at least two of the gRNAs, expression of *CEP250* and *CPNE1* were diminished relative to control cells, which received the dCas9 construct without an accompanying gRNA (**Fig 5B**), suggesting that dCas9/gRNA complexes had obstructed binding or function of a transcriptional enhancer at this site. This result is directionally consistent with the data from the isogenic HUES 8 rs2277862 deletion mutants, which also indicated that the major allele (C) has enhancer activity, though at odds with the MPRA data that suggested the minor allele (T) has enhancer activity.

**Fig. 5.**
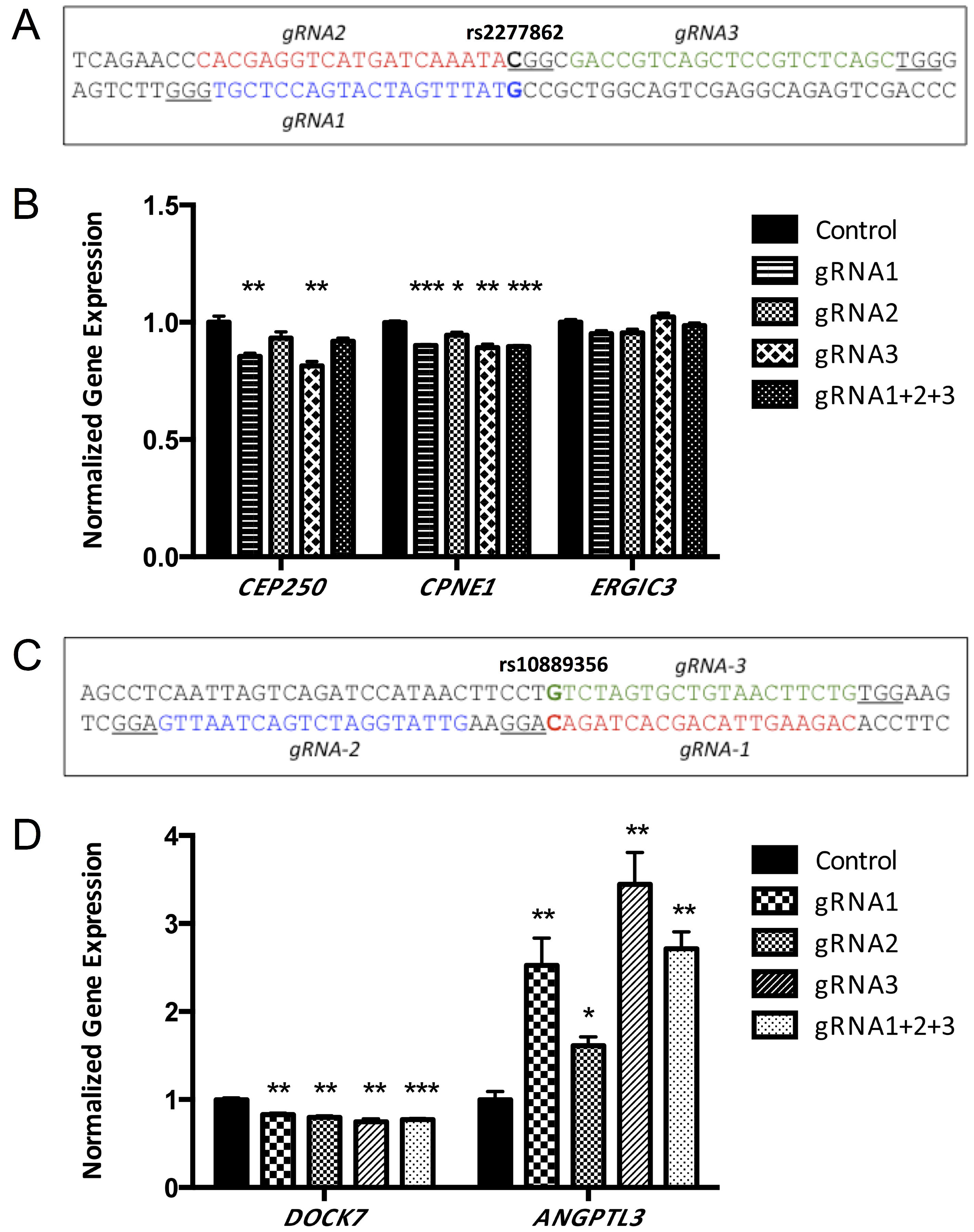
CRISPR interference modulates gene expression at the 20q11 and 1p31 loci. (A)Guide RNAs used at the rs2277862 site. The guide RNA protospacers are in colors, the PAMs are underlined, and the SNP position is indicated in bold. (B) Gene expression, normalized to mean expression level in control cells, in HEK 293T cells (homozygous major at rs2277862) transfected with catalytically dead Cas9 (dCas9) with various gRNAs targeting the rs2277862 site (either singly or in combination, 3 wells per condition). Control cells were transfected with dCas9 without an accompanying gRNA. (C) Guide RNAs used at the rs10889356 site. (D) Gene expression in HepG2 hepatoma cells (homozygous major at rs10889356) transfected with dCas9 with various gRNAs targeting the rs10889356 site (3 wells per condition). Data are displayed as means and s.e.m. (^*^*P*<0.05, ^**^*P*<0.01, ^***^*P*<0.001 relative to control, calculated by two-tailed unpaired Student’s *t*-test).

We validated rs10889356 as a causal variant with CRISPRi constructs targeting the rs10889356 site in HepG2 cultured hepatoma cells, which are homozygous major (G/G) at this SNP (**Fig 5C**). We chose HepG2 cells so that we could assess not just *DOCK7*but also *ANGPTL3*, the latter being liver-specific. In light of the low transfection efficiency of HepG2 cells, positive transfectants were isolated by fluorescence-activated cell sorting (FACS) prior to gene expression analysis. With three different gRNAs, singly or in combination, expression of *DOCK7* was significantly decreased and expression of *ANGPTL3* was significantly increased relative to control cells, which received the dCas9 construct without an accompanying gRNA (**Fig 5D**). These results are directionally consistent with the data from the isogenic H7 HLC rs10889356 deletion mutant experiment, which was also performed on a homozygous major background. Additionally, both experiments recapitulated the inverse relationship between *DOCK7* and *ANGPTL3* expression levels revealed by the human eQTL data for the locus lead SNP rs2131925 [2].

### Interrogation of an rs2277862 knock-in mouse

Because independent hPSC clones displayed variable propensities for directed differentiation, thereby introducing confounding variability in gene expression studies, we sought an alternative approach to faithfully model the effect of regulatory variation in primary tissues of interest, namely liver and fat. The noncoding region encompassing rs2277862 is well conserved in mouse (**Fig 6A**). Remarkably, the orthologous nucleotide in mouse also displays naturally occurring variation and has been previously cataloged as rs27324996 on chromosome 2, with the same two alleles (C and T) as in humans. As documented in dbSNP, rs27324996 has a MAF of 43% based on genotyping analyses of 14 different inbred strains of mice. All three human eQTL genes have a murine ortholog at this locus, and the orientation of these genes relative to the putative regulatory variant is conserved between mouse and human (**Fig 6A**).

**Fig. 6.**
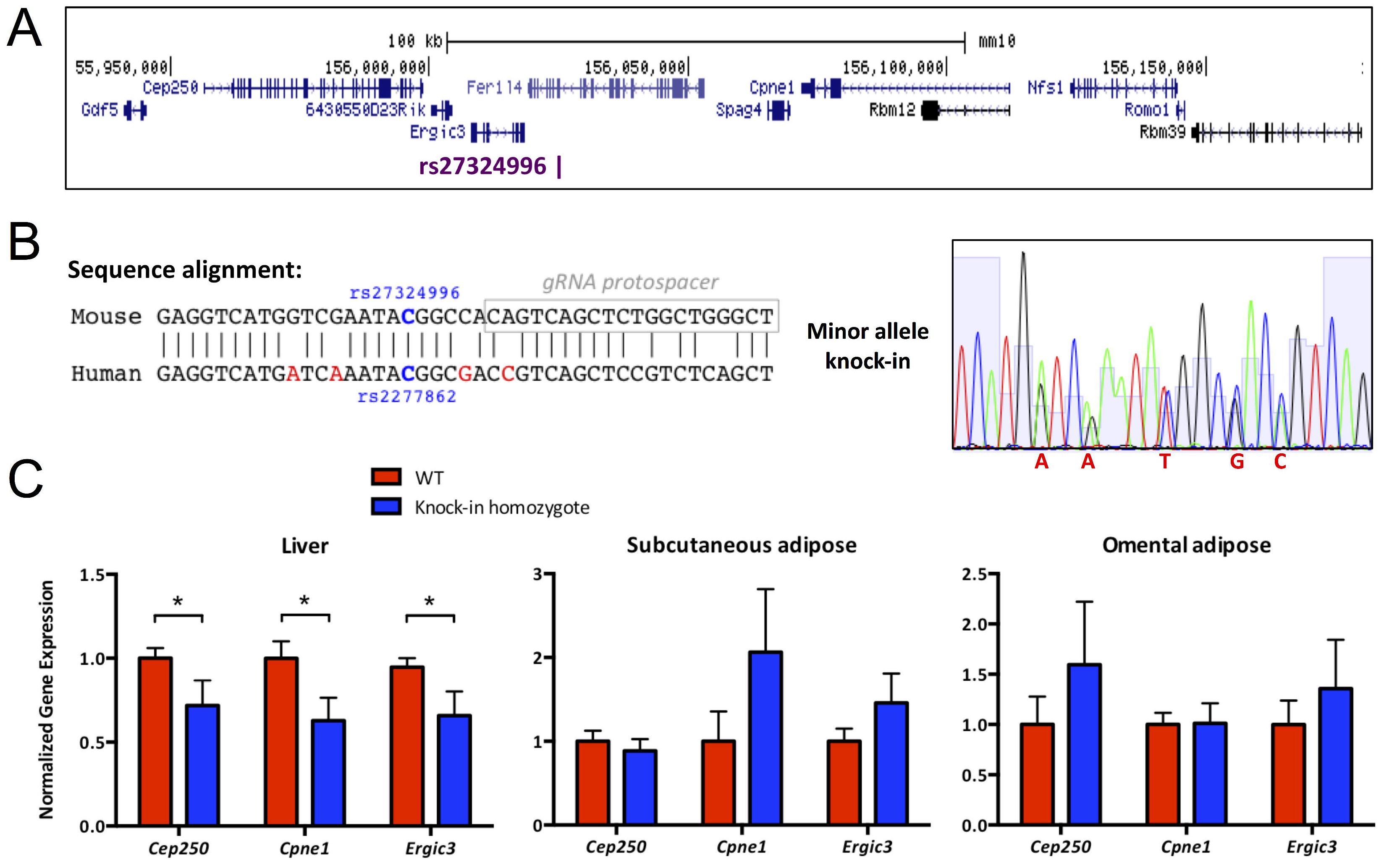
Gene expression in an rs2277862 knock-in mouse. A) Schematic of mouse chromosome 2qH1 locus showing the relative positions of rs27324996, *Cep250, Cpnel*, and *Ergic3*. The architecture closely matches the orthologous human chromosome 20q11 locus. (B) The noncoding region encompassing rs2277862 is well conserved in mouse, including allelic variants of the SNP itself, with the murine equivalent being rs27324996. The SNP position is indicated in blue bold, non-conserved nucleotides are indicated in red, and the guide RNA protospacer used to generate the knock-in mouse is boxed. The electropherogram is from a mouse in which the minor allele of rs2277862/rs27324996 (T) has been knocked into one chromosome, along with the four non-conserved nucleotides that “humanize” the site. (C) Gene expression in liver and fat tissues from littermates of the C57BL/6J background (*N* = 18 wild-type mice and 10 homozygous knock-in mice), normalized to mean expression level in wild-type mice. Data are displayed as means and s.e.m. (^*^*P*<0.05, ^**^*P*<0.01, ^***^*P*<0.001 relative to control, calculated by two-tailed unpaired Student’s *t*-test).

Because MPRA identified a transcriptional role for rs2277862 when performed in 3T3-L1 adipocytes, which are of murine origin, we reasoned that the regulatory machinery at this site may be present in mouse tissues *in vivo*. Since HDR is far more efficient in mouse embryos than in hPSCs [16], we used CRISPR-Cas9 to generate a knock-in mouse with a minor allele on the C57BL/6J background, which is homozygous major (C/C) at rs27324996. Compared to genome editing of hPSCs, one clear advantage of a mouse model is that even if only a single positive founder mouse is obtained with genome editing, the knock-in allele can be bred to homozygosity in a matter of months, and a large number of wild-type and knock-in mice can be generated for well-powered gene expression studies in liver, subcutaneous fat, and omental fat. In principle, this study design could enable replication of human eQTL association data while avoiding the disadvantages of cultured human transformed cell lines (such as HepG2) and of suboptimal and variable hPSC differentiation.

One-cell mouse embryos were injected with Cas9 mRNA, a gRNA targeting the site of rs27324996, and a ssDNA donor template carrying the minor SNP allele as well as four additional nucleotide changes that served to both “humanize” the sequence (i.e., make a perfect match to the orthologous human sequence) and prevent re-cleavage of the knock-in allele by CRISPR-Cas9. Out of 37 founder mice, there was one mouse bearing the knock-in allele (**Fig 6B**). We bred this mouse through two generations to obtain homozygous minor allele knock-in mice.

We compared *Cep250, Cpne1*, and *Ergic3*gene expression in primary liver, subcutaneous fat, and omental fat from wild-type (C/C) and homozygous knock-in (T/T) C57BL/6J littermates (*n* = 18 wild-type and 10 knock-in mice). In liver, there was significantly decreased expression of *Cep250* (down 28.2%), *Cpne1* (down 37.2%) and *Ergic3* (down 34.1%) in the homozygous knock-in mice (**Fig 6C**). Notably, these changes are quite concordant with the directionality and magnitudes of the changes observed between wild-type (C/C) and homozygous knockout hPSCs (compare with **Fig 3B**). In fat tissues from knock-in mice, no statistically significant differences were apparent, due to a large degree of variance in the gene expression measurements (**Fig 6C**). Although no conclusions can be drawn from the fat data, the liver data from the knock-in mice in combination with the genome-edited hPSC data and the CRISPRi data suggest that the site of rs2277862/rs27324996 has transcriptional regulatory activity in both human and mouse.

## Discussion

One of the principal challenges of the post-GWAS era has been cataloging the allelic spectrum of causal variants underlying complex trait susceptibility. In this work, we describe a methodological framework for causal variant discovery that involves high-throughput identification of putative disease-causal loci through a functional reporter-based screen, MPRA, followed by validation of prioritized variants in genome-edited cellular models. As a proof-of-concept, we focused our experimental efforts on validating two top-ranked MPRA variants that are potentially causal for lipid phenotypes.

We sought to rigorously demonstrate causality for rs2277862 and rs10889356 through the generation of isogenic wild-type and mutant hPSCs. Consistent with prior studies, we found the generation of knock-in clones via HDR to be very inefficient compared to the generation of clones with defined deletions via multiplexed NHEJ. The difference in efficiency proved to be crucial to the relative success of the alternative study designs. We were only able to generate a single rs2277862 heterozygous knock-in clone despite screening hundreds of clones. We found that clone-to-clone variability was sufficiently high to swamp out the expected small differences in gene expression—on the order of 10% to 20% differences, based on eQTLs from human tissue studies—and, accordingly, little informative data was obtained from the knock-in hPSC clone. The situation was even worse when using cells differentiated from the hPSCs, given the increased clone-to-clone variability that resulted.

It was feasible to obtain more precise data only when numerous wild-type and mutant clones were used in the experiments (i.e., large n), which in turn was only possible when using NHEJ to generate knockout clones with high efficiency. The knockout study design proved fruitful for both rs2277862 and rs10889356, with knockout clones displaying statistically significant, albeit small, differences from wild-type clones. Even with the use of a multitude of clones, however, differentiation of the hPSCs greatly increased the clone-to-clone variability and obscured any differences between wild-type and knockout clones.

This is not to say that hPSCs are a poor platform for functional genetic studies. hPSCs offer many attractive advantages over existing model systems, including genomic stability, pluripotency, and renewability. However, it seems prudent to consider the estimated “signal” from a putative disease-causal variant relative to the estimated “noise” from a directed differentiation protocol prior to pursuing a disease-modeling experiment in hPSCs. For highly penetrant variants with large effect sizes, hPSCs have been shown to yield informative insights into human genetics. For example, Ding et al. were able to characterize the rare gain-of-function variant E17K in the gene *AKT2*, which is associated with hypoglycemia, hypoinsulinemia, and increased body fat secondary to dysregulated insulin signaling [17]. hPSC-derived *AKT2* E17K knock-in HLCs displayed significantly decreased glucose production, while hPSC-derived adipocytes had increased TG content and increased glucose uptake compared to matched wild-type controls. Through disease modeling in hPSCs, Ding et al. unequivocally established a dominant activating role for the *AKT2* E17K mutation. However, GWAS-implicated common variants would be expected to display much smaller effect sizes, suggesting that hPSC-derived cell types may not be the ideal model system in which to investigate common genetic variation, despite the touted benefits of isogenic disease modeling.

We explored two alternative model systems. The first involved CRISPR interference, a term that is typically used to describe the repression of transcription via the positioning of dCas9 to a gene promoter or coding sequence to block transcriptional initiation or elongation via steric interference [15]; alternatively, dCas9 can be attached to a repressor domain such as Krüppel associated box (KRAB) to effect transcriptional silencing via epigenetic chromatin modification [18]. We opted to use the unadorned dCas9 protein rather than attaching an extra domain, since the regulatory elements in the vicinity of the two SNPs are undefined and it was unclear whether each SNP allele acted as an enhancer or repressor or was neutral. Instead, we relied on steric interference to reverse any effect of the SNP allele. For both rs2277862 and rs10889356, CRISPRi yielded results that were concordant with the effects observed in knockout hPSCs and differentiated cells—decreased gene expression vis-à-vis the major allele of rs2277862, and decreased *DOCK7* and increased *ANGPTL3* expression vis-à-vis the major allele of rs10889356.

Of note, we performed the CRISPRi experiments in cultured human transformed cells rather than hPSCs, making the experiments far easier to perform and, in the case of HepG2 cells, allowing us to use cells with inherent hepatocyte properties without having to undertake the differentiation of hPSCs into HLCs. Indeed, our findings suggest that CRISPRi may be better suited to the high-throughput interrogation of candidate causal variants (whether nominated by MPRA or another technique) than genome editing of cells, although the latter should be regarded as the gold standard, especially when it entails the knock-in of a specific allelic variant. Furthermore, a limitation of CRISPRi is that it relies on the major allele of a SNP having enhancer or repressor
activity (rather than being neutral), since it may not always be possible to identify commonly used human cultured transformed cell lines with minor alleles.

The second alternative model system we attempted to use was a knock-in mouse. This is not a generalizable strategy, as it depends on the human SNP sequence and the orthologous sequence in the mouse genome being very closely matched, which is not often the case with noncoding regions. (One possible means to overcome this limitation would be the introduction of the entirety of the orthologous human locus into the mouse genome, via bacterial artificial chromosome transgenesis or another approach.) Fortuitously, the human rs2277862 sequence matches closely with mouse, to the extent that the mouse genome has a perfectly corresponding SNP, rs27324996. This allowed us to generate a “humanized” minor allele homozygous mouse to compare with the wild-type major allele homozygous mouse. The theoretical advantage of this study design is that it allows for the analysis of authentic primary tissues such as liver and fat without the need for directed differentiation *in vitro* or reliance on transformed cell lines. In practice, while we observed statistically significant trends in mouse liver that were concordant with our findings in genome-edited hPSCs, we found that there was substantial mouse-to-mouse variability with respect to gene expression of *Cep250, Cpne1*, and *Ergic3* in fat. Thus, the challenges inherent in studying common DNA variants with small effect sizes may apply to both mice and hPSC models. One difference between the model systems, of course, is that it would be much easier to generate a very large number of isogenic mice, through breeding, than it would be generate a very large number of independent isogenic hPSC clones in order to increase the power of a study.

Finally, we note the discordance among some of the results obtained from the MPRA experiments and the genome editing/CRISPRi experiments. While the results for rs10889356 were concordant—all three types of experiments agreed that the major allele (G) has enhancer activity—with respect to rs2277862, the MPRA data suggest that the minor allele (T) has more enhancer activity on a closely linked reporter gene, while the genome editing/CRISPRi data suggest that the major allele (C) has more enhancer activity on a more distant endogenous gene. A salient difference between the two types of experiments is that the MPRA experiments assessed heterologously expressed 144-nt DNA fragments shorn of their genomic context, which could substantially alter their effects on gene expression. In contrast, both the genome editing and CRISPRi experiments targeted endogenous sequences. In light of this distinction, MPRA experiments should be considered exploratory only, whereas genome editing and CRISPRi experiments should be considered more representative of regulatory effects in endogenous loci, and accordingly, requisite confirmation for any results obtained by MPRAs.

In summary, these studies have highlighted the utility of high-throughput functional genomics approaches to prioritize putative causal GWAS variants, and they provide validation of rs2277862 and rs10889356 as casual eQTL variants for two lipid-associated loci. These studies also illustrate the challenges inherent in modeling common genetic variation in available model systems.

## Materials and Methods

### Massively parallel reporter assays

MPRA experiments were performed as previously described [10]. Approximately 240,000 144-bp oligonucleotides representing ~11,000 distinct “tiles” with the major or minor alleles of 1,837 candidate causal variants in the center, left-shifted, or right-shifted positions and coupled to distinguishing barcodes were generated by microarray-based DNA synthesis (Agilent). The tiles and barcodes were separated by two common restriction sites. The oligonucleotides were PCR amplified using universal primer sites and directionally cloned into a pMPRA1 (Addgene plasmid #49349) backbone using Gibson assembly. A minimal promoter-firefly luciferase segment from pMPRAdonor2 (Addgene plasmid #49353) was inserted between the tiles and barcodes via double digestion and directional ligation. The resulting reporter plasmid pools were co-transfected into either undifferentiated 3T3-L1 cells (pre-adipocytes) or differentiated 3T3-L1 adipocytes using FuGENE 6 (Promega). Two biological replicate MPRA experiments were performed in each cell type. The relative enhancer activities of the different tiles were calculated by comparing the corresponding barcodes from the cellular mRNA and the transfected plasmid pool. The highest-priority “hits” were tiles for which there was the greatest difference (in magnitude and statistical significance) in enhancer activity between the major and minor alleles in multiple positions (center, left-shifted, right-shifted) in replicate experiments in both adipocytes and pre-adipocytes.

For the single-hit saturation mutagenesis MPRA experiments [10], all possible single substitutions were introduced at all positions (expect the position of the SNP) within the rs2277862 right-shifted tile (with either the major or minor allele) or the rs10889356 left-shifted tile (with either the major or minor allele). The pools of variant tiles were generated, introduced into reporter constructs, transfected into 3T3-L1 adipocytes, and analyzed as described above. The plots in **Fig 1** were generated as previously described [10].

### CRISPR-Cas9 plasmid construction

Guide RNAs were designed by manual inspection of the genomic sequence flanking rs2277862 and rs10889356 and evaluated for potential off-target activity using the CRISPR Design tool at http://crispr.mit.edu. Protospacers were cloned into the BbsI site of pGuide (Addgene plasmid #64711) via the oligonucleotide annealing method, and, if not already present, a G was added to the 5’ end to facilitate U6 polymerase transcription. Genome editing was performed using pCas9_GFP (Addgene plasmid #44719), which co-expresses a human codon-optimized Cas9 nuclease and GFP via a viral 2A sequence.

For CRISPRi studies, pAC154-dual-dCas9VP160-sgExpression (Dr. Rudolph Jaenisch, Addgene plasmid #48240), a dual expression construct that expresses dCas9-VP160 and sgRNA from separate promoters, was modified by PCR-based methods to remove the VP160 domain and include a viral 2A sequence and GFP after dCas9. Additionally, the gRNA sequence was modified to include a 5-bp hairpin extension, which improves Cas9-gRNA interaction, and a single base pair substitution (A-U flip) that removes a putative Pol III terminator sequence, as described previously [19].

### Cultured cell line maintenance and transfection

All cell lines were maintained in a humidified 37°C incubator with 5% CO2. HEK 293T, HepG2, and 3T3-L1 cells were cultured in high glucose DMEM supplemented with 10% FBS and 1% penicillin/streptomycin. For MPRA experiments, 3T3-L1 cells were differentiated into adipocytes with the addition of 0.5 mM IBMX, 1μM dexamethasone, and 10 μg/mL insulin to the media for 3 days, followed by addition of only 10 μg/mL insulin for another 3 days. For CRISPRi experiments, HEK 293T and HepG2 cells were seeded into 6-well plates and transfected 24 hours later using Lipofectamine 3000 (Life Technologies) according to the manufacturer’s instructions.

### hPSC culture and CRISPR-Cas9 targeting

HUES 8 and H7 cells were grown under feeder-free conditions on Geltrex (Life Technologies)-coated plates in chemically defined mTeSR1 medium (STEMCELL Technologies) supplemented with 1% penicillin/streptomycin and 5 μg/mL Plasmocin (InvivoGen). Medium was changed every 24 hours. For electroporation, cells in a 60%-70% confluent 10-cm plate were dissociated into single cells with Accutase (Life Technologies), resuspended in PBS, and combined with 25 Hg pCas9_GFP and 25 μg gRNA plasmid (or 12.5 μg of two different gRNA plasmids, for multiplexed targeting) in a 0.4 cm cuvette. For knock-in, 15 μg pCas9_GFP, 15 ng gRNA plasmid, and 30 ng ssODN (5’-GGTCGTCAGAACCCACGAGGTCATGATCAAATATGGCGACCGTCAGCTCCGTCTCAGCTGGGAGAGA-3’) were used instead. A single pulse was delivered at 250 V/500 μF (Bio-Rad Gene Pulser), and the cells were recovered and plated in mTeSR1 with 0.4 μM ROCK inhibitor (Y-27632, Cayman Chemical). Cells were dissociated with Accutase 48 hours postelectroporation, and GFP-positive cells were isolated by FACS (FACSAriaII, BD Biosciences) and replated onto 10-cm Geltrex-coated plates (15,000 cells/plate) with conditioned medium and 0.4 μM ROCK inhibitor to facilitate recovery.

### Isolation and screening of clonal hPSC populations

Following FACS, single cells were permitted to expand for 10-14 days to establish clonal populations. Colonies were manually picked and replated into individual wells of a 96-well plate. Once the wells reached 80-90% confluence, cells were dissociated with Accutase and split at a 1:3 ratio to create a frozen stock and two working stocks that were maintained in culture. For genomic DNA isolation, cells from one of the working stocks were lysed in 50 μL lysis buffer (10 mM Tris pH 7.5, 10 mM EDTA, 10 mM NaCl, 0.5% Sarcosyl) with 40 μg/mL Proteinase K for 1-2 hours in a humidified incubator at 56°C. Genomic DNA was precipitated by addition of 100 μL 95% ethanol with 75 mM NaCl, followed by incubation at −20°C for 2 hours. Precipitated DNA was washed three times with 70% ethanol, resuspended in 30-50 μL TE buffer with 0.1 mg/mL RNase A, and allowed to dissolve at room temperature overnight.

HPSC clones were screened by PCR amplification of a small region surrounding the targeted site using BioReady rTaq DNA Polymerase (Bulldog Bio) and the following cycling conditions: 94°C for 5 min, [94°C for 30 sec, 54-56.5°C for 30 sec, 72°C for 30 sec] × 40 cycles, 72°C for 5 min. The following primer pairs were used: for rs2277862, F: 5’-TGCTGGACCCACACTTCATA-3’ and R: 5’-CTCAGTCCCTCTCCCTCCTT-3’; for rs10889356, F: 5’-CCATTAGGTCACTTGCCAGA-3’ and R: 5’-ACAGGGGGATTCTGTCTAAAA-3’. PCR amplicons were separated on a high-percentage agarose gel, and clones with indels were identified based on size shifts relative to the wild-type band. Suspected mutant clones were confirmed by Sanger sequencing of the PCR products.

Multiple mutant clones were retrieved from the frozen stock, or if possible, from the second working stock and expanded for experiments. Additionally, several clones that underwent the targeting procedure but remained genetically wild-type at the intended site were expanded as controls.

### Differentiation of hPSCs into white adipocytes

Differentiation of HUES 8 cells to white adipocytes was performed as previously described [20]. To induce embryoid body formation, wild-type and mutant hPSCs were pre-treated overnight with 2% DMSO; dissociated into small clumps with Accutase; resuspended in growth medium containing DMEM, 10% knockout serum replacement (Life Technologies), 2 mM GlutaMAX (Life Technologies), 1% non-essential amino acids, 1% penicillin/streptomycin, and 0.1 mM beta-mercaptoethanol; and transferred to low-attachment 6-well plates (Costar Ultra Low Attachment; Corning Life Sciences). After one week in culture, embryoid bodies were collected and replated onto gelatin-coated plates in MPC medium containing DMEM, 10% FBS, 1% penicillin/streptomycin, and 2.5 ng/mL bFGF (Aldevron). Cells were serially passaged at a 1:3 ratio to obtain a homogenous population of MPCs by passage 3-4.

Recombinant lentivirus was produced using a third-generation, Tat-free packaging system. Lentiviral vectors encoding either doxycycline-inducible *PPARG2* or rtTA were transfected into HEK 293T cells by the calcium phosphate method, along with the packaging plasmids pMDL and pREV and a capsid plasmid encoding VSV-G. Viral supernatant was harvested at 48 and 72 hours post-transfection and filtered through a 0.45 μm membrane. One day before transduction, MPCs were plated at 1 × 10^6^ cells per 10-cm plate. The following day, MPCs were transduced with 5 mL lenti-*PPARG2* and 5 mL lenti-rtTA and incubated at 37°C for 16 hours. After the viral supernatant was aspirated, the cells were washed with PBS and allowed to grow to confluence. Transduced MPCs were split into 6-well dishes prior to initiating white adipocyte differentiation.

Differentiation was induced by the addition of adipogenic media containing DMEM, 7.5% knockout serum replacement, 7.5% human plasmanate (Grifols), 0.5% non-essential amino acids, 1% penicillin/streptomycin, 0.1 μM dexamethasone (Sigma), 10 μg/mL insulin (Sigma), and 0.5 μM rosiglitazone (Santa Cruz). The differentiation medium was supplemented with 700 ng/mL doxycycline from day 0 to 16. Doxycycline was then removed from the culture medium until day 21, at which point the differentiated cells were used for experiments.

### Differentiation of hPSCs into hepatocyte-like cells

Differentiation of H7 cells into HLCs was performed as previously described [17]. One day before differentiation, hPSCs at 60% confluence were split at a 1:3 ratio into 6-well dishes with mTeSR1 plus 0.4 μM ROCK inhibitor. Cells were serially cultured in (1) RPMI-B27 (RPMI-1640 from Sigma; B27 supplement minus Vitamin A from Life Technologies) supplemented with 100 ng/mL Activin A (PeproTech) and 3μM CHIR99021 (Cayman Chemical), a glycogen synthase kinase 3 inhibitor, for 3 days to obtain definite endoderm; (2) RPMI-B27 supplemented with 5 ng/mL bFGF (Millipore), 20 ng/mL BMP4 (PeproTech), and 0.5% DMSO for 5 days to obtain hepatic endoderm; (3) RPMI-B27 supplemented with 20 ng/mL HGF (PeproTech) and 0. 5% DMSO for 5 days to obtain hepatoblasts; and (4) Hepatocyte Culture Medium (Lonza) supplemented with 20 ng/mL HGF, 20 ng/mL Oncostatin M (PeproTech), 100 nM dexamethasone (Sigma), and 0.5% DMSO for 10 to 12 days to obtain HLCs.

### CRISPR knock-in mice

Four candidate guide RNAs with a cut site near rs27324996 were designed by manual inspection, and the corresponding protospacers were cloned into the pGuide plasmid as described above. Each gRNA plasmid was co-transfected with pCas9_GFP into mouse 3T3-L1 cells using TransIT-2020 Reagent (Mirus Bio) according to the manufacturer’s instructions. Two days posttransfection, GFP-positive cells were isolated by FACS, and genomic DNA was isolated using the DNeasy Blood and Tissue Kit (Qiagen). The region flanking rs27324996 was PCR amplified (F: 5’-TGGGAATGGCTTCTTAGGGC-3’ and R: 5’-CATCCCCAAGCAACTCAACC-3’) using AccuPrime Taq DNA Polymerase (Life Technologies) with the following cycling conditions: 94°C for 2 min, [94°C for 30 sec, 55°C for 30 sec, 68°C for 30 sec] × 40 cycles, 68°C for 5 min. PCR products were purified using the DNA Clean and Concentrator kit (Zymo Research) and analyzed for the presence of indels using the Surveyor Mutation Detection Kit (IDT) according to the manufacturer’s instructions. Cel-I nuclease-treated PCR products were resolved on a 1.5% agarose gel to detect mutagenesis activity. The gRNA sequence exhibiting the highest mutation rate was PCR amplified, and the purified PCR product was used as a template for *in vitro* transcription using the MEGAshortscript T7 kit (Life Technologies). The transcribed RNA was purified by phenol/chloroform extraction, ethanol precipitated, and resuspended in injection buffer (5 mM Tris-HCl pH 7.6, 0.1 mM EDTA).

All animal procedures described here were reviewed and approved by the Harvard University Institutional Animal Care and Use Committee (protocol #14-05-202). Euthanasia in all instances was via terminal inhalation of carbon dioxide, consistent with the 2013 AVMA Guidelines on Euthanasia.

One-cell embryo injections were performed by the Genome Modification Facility at Harvard University. Superovulated C57BL/6J females were mated with C57BL/6J males, and fertilized embryos were harvested from the oviducts. One-cell embryos were injected with a mixture of 100 ng/μL Cas9 mRNA (TriLink BioTechnologies), 50 ng/μL gRNA, and 100 ng/μL ssODN (5’-AGCCCACAGTTGGCTCTGTGGTGGCTATAGAATCTGTTTTCCAGGTCAATGTG GGTCTCCCCGATGAGGTCATCTGAACCCACGAGGTCATGATCAAATATGGCG ACCGTCAGCTCTGGCTGGGCTGGGAGGGAGACGCTCAGCTCCAGGACCCTGG GCAGGAAGGGAAATTGACTAACCACAGCTCCATGCCCTCAGAG-3’). Injected embryos were implanted into the uteri of pseudopregnant foster mothers.

DNA was prepared from tail biopsies of 3-week-old founder mice by the hot hydroxide method, and genotyping was performed with the same PCR primers and cycling conditions used for the Cel-I nuclease assay. Positive founders were identified by Sanger sequencing of PCR products. The single positive founder was bred to a wild-type C57BL/6J mouse (Jackson Laboratories), and the resulting progeny were intercrossed for one to two generations to breed the knock-in allele to homozygosity. Genotyping of progeny was performed in the same manner. Wild-type and homozygous knock-in littermates from several litters, ~12 weeks of age, were used for gene expression analyses.

### Quantitative reverse transcriptase-polymerase chain reaction

Wells of cells were washed with ice-cold PBS and lysed directly in TRIzol Reagent (Thermo Fisher Scientific), and primary liver and fat samples from mice were homogenized in TRIzol Reagent. RNA was isolated according to the manufacturer’s instructions and reverse transcribed using SuperScript III Reverse Transcriptase (Thermo Fisher Scientific) with an equimolar mixture of random hexamers and oligo-dT. Gene expression was measured using the following TaqMan assays (Applied Biosystems): Hs00898245_m1 for *CEP250*, Hs00537765_m1 for *CPNE1*, Hs00211070_m1 for *ERGIC3*, Hs00205581_m1 for *ANGPTL3*, Hs00290630_m1 for *DOCK7*, Hs00910225_m1 for *ALB*, Hs01097800_m1 for *SERPINA1*, Mm00623502_m1 for *Cep250*, Mm00467970_m1 for *Cpne1*, and Mm00499400_m1 for *Ergic3*. Human *B2M*(Assay ID 4326319E) or mouse *Actb* (Assay ID 4352341E) was used as the reference gene. Each 10 μL qPCR reaction contained 1 μL cDNA (diluted 1:3 with water) and was performed in technical duplicate or triplicate. Reactions were carried out on a ViiA 7 Real-Time PCR system (Applied Biosystems), and relative expression differences were quantitated by the ΔΔC_t_ method.

## Acknowledgments

We are grateful to Qiurong Ding, Alexandra Chadwick, and Alanna Strong for critical reading of the manuscript.

## Supplemental Information

### Supporting Information Captions

**S1 Table. Candidate SNPs in adipose lipid-associated eQTLs that were interrogated in the MPRA experiments**.

**S2 Table. MPRA experimental data**. Each row corresponds to a 144-bp tile either centered (C), right-shifted (R), or left-shifted (L) relative to a SNP, with the two alleles indicated by the last two letters in the label for each row (penultimate is allele 1, last is allele 2). Each allele on each tile was repeated in the design with 22 different barcodes, although due to synthesis and cloning biases, some have too many dropouts to give robust results (marked by NaN). Four independent transfection experiments were performed: L1Ad_R1 (3T3-L1 adipocytes #), L1Ad_R2 (adipocytes #2), L1Pre_R1 (3T3-L1 pre-adipocytes #1), and L1Pre_R2 (pre-adipocytes #2). For each SNP and transfection there are 6 columns: ^*^_Sig1 = signal from allele 1 (log of median barcode counts for this tile divided by median barcode counts for all tiles, with a high positive value meaning enhancer activity, and a negative value meaning repressor/silencer activity); ^*^_Sig1_P = Mann-Whitney U-test P-value for the signal from allele 1 being the same as the median signal from all tiles (not corrected for multiple testing); ^*^_Sig2 = signal from allele 2; ^*^_Sig2_P = P-value for allele 2; ^*^_Var = log-ratio of signal from allele 1 over signal from allele 2; ^*^_Var_P = P-value for the signal from alleles 1 and 2 being the same.

